# Development of avian influenza A(H5) virus datasets for Nextclade enables rapid and accurate clade assignment

**DOI:** 10.1101/2025.01.07.631789

**Authors:** Jordan T. Ort, Sonja A. Zolnoski, Tommy T.-Y. Lam, Richard Neher, Louise H. Moncla

## Abstract

The ongoing panzootic of highly pathogenic avian influenza (HPAI) A(H5) viruses is the largest in history, with unprecedented transmission to multiple mammalian species. Avian influenza A viruses of the H5 subtype circulate globally among birds and are classified into distinct clades based on their hemagglutinin (HA) genetic sequences. Thus, the ability to accurately and rapidly assign clades to newly sequenced isolates is key to surveillance and outbreak response. Co-circulation of endemic, low pathogenic avian influenza (LPAI) A(H5) lineages in North American and European wild birds necessitates the ability to rapidly and accurately distinguish between infections arising from these lineages and epizootic HPAI A(H5) viruses. However, currently available clade assignment tools are limited and often require command line expertise, hindering their utility for public health surveillance labs. To address this gap, we have developed datasets to enable A(H5) clade assignments with Nextclade, a drag-and-drop tool originally developed for SARS-CoV-2 genetic clade classification. Using annotated reference datasets for all historical A(H5) clades, clade 2.3.2.1 descendants, and clade 2.3.4.4 descendants provided by the Food and Agriculture Organization/World Health Organization/World Organisation for Animal Health (FAO/WHO/WOAH) H5 Working Group, we identified clade-defining mutations for every established clade to enable tree-based clade assignment. We then created three Nextclade datasets which can be used to assign clades to A(H5) HA sequences and call mutations relative to reference strains through a drag-and-drop interface. Nextclade assignments were benchmarked with 19,834 unique sequences not in the reference set using a pre-released version of LABEL, a well-validated and widely used command line software. Prospective assignment of new sequences with Nextclade and LABEL produced very well-matched assignments (match rates of 97.8% and 99.1% for the 2.3.2.1 and 2.3.4.4 datasets, respectively). The all-clades dataset also performed well (94.8% match rate) and correctly distinguished between all HPAI and LPAI strains. This tool additionally allows for the identification of polybasic cleavage site sequences and potential N-linked glycosylation sites. These datasets therefore provide an alternative, rapid method to accurately assign clades to new A(H5) HA sequences, with the benefit of an easy-to-use browser interface.

## Introduction

Highly pathogenic avian influenza (HPAI) A viruses of the H5 subtype continue to pose risks to human and animal health, with transmission in wild and domestic birds seeding periodic cross-species transmission and outbreaks [1, 2]. First detected in China in 1996 with the A/Goose/Guangdong/1/1996 (GsGd) strain, these viruses have now spread worldwide and are currently causing the largest panzootic of HPAI in history [3, 4]. Highly pathogenic A(H5Nx) viruses—those carrying an H5 hemagglutinin (HA) and various subtypes of NA (e.g. N1, N2, N3, N5, N6, and N8)—have greatly diversified since the emergence of the GsGd lineage, a result of both substitutions in HA and reassortment with other gene segments [5]. Through this accumulation of HA mutations, A(H5Nx) viruses have evolved into distinct phylogenetic groups, known as clades, based on their HA sequence [6].

A(H5) clades are defined by the Food and Agriculture Organization/World Health Organization/World Organisation for Animal Health (FAO/WHO/WOAH) H5 Working Group, who periodically reviews global A(H5) virus genetic sequence data to determine whether there is sufficient intra-clade diversity to support designating sublineages as new clades [7–11]. Under this nomenclature system, clades are defined as monophyletic groups having <1.5% within-clade average pairwise nucleotide distance, >1.5% distance between other clades, and at least 60% bootstrap support [7]. Currently, high levels of circulating diversity within the 2.3.2.1c and 2.3.4.4 clades necessitate the division of each, yielding subclades 2.3.2.1d–g and 2.3.4.4a–h. In addition to these HPAI clades, there are two lineages of primarily low pathogenic avian influenza (LPAI) viruses that remain endemic in wild birds and are denoted American and Eurasian non-GsGd (Am-nonGsGd and EA-nonGsGd, respectively). This nomenclature system allows for consistent classification and interpretation of surveillance data across laboratories [7]. The ability to classify A(H5) HA sequences provides a framework for studying A(H5Nx) virus evolution, including changes in antigenicity or pathogenicity, host adaptation, and reassortment [7, 12–14]. By enabling researchers to rapidly distinguish the frequency of lineages that are currently circulating among birds and other animals in different regions, this system is critical to viral surveillance and public health monitoring. Clade classifications are therefore crucial for accurately tracking the spread of A(H5Nx) viruses, identifying outbreaks, and improving public health outcomes through more informed decision-making and risk assessment.

Despite the importance of this nomenclature system, A(H5) clade annotation tools are limited. One popular tool for A(H5) clade assignment is LABEL [15] (https://wonder.cdc.gov/amd/flu/label/), which is a command-line software that utilizes a hidden Markov model to classify HA sequences without an alignment step. While using an alignment-free approach allows for rapid clade assignments, this methodology obscures the ability to identify amino acid substitutions and key features of HA—such as the cleavage site sequence—and to assess sequence quality based on the presence of indicators like frameshifts, indels, and premature stop codons. Additionally, LABEL is a command-line software without a graphical user interface, posing a barrier to researchers in public health and surveillance laboratories without bioinformatics expertise or command line access. Alternative web-based tools are available, such as those from the Bacterial and Viral Bioinformatics Resource Center (BV-BRC; https://www.bv-brc.org) and The University of Hong Kong (https://tipars.hku.hk/reference/Influenza-A-H5_HA_Tree), which can classify A(H5) HA sequences into clades and perform phylogenetic placement [16]. However, these cannot be used to assess sequence quality, identify amino acid motifs, or call mutations relative to reference strains.

To address these gaps, we have developed datasets to allow for A(H5) clade assignment with Nextclade, a drag-and-drop browser tool originally developed for SARS-CoV-2 classification that can be run via the command line or via a web interface [17]. Each Nextclade dataset is composed of a reference phylogeny in which each tip and internal node has been annotated with a clade assignment. To perform clade assignments, Nextclade first aligns and trims each query sequence relative to the dataset-specific reference sequence. Next, mutations in each query sequence are compared to the sets of mutations possessed by each node and tip on the tree. After identifying the node or tip with the most similar set of mutations (see https://docs.nextstrain.org/projects/nextclade/en/stable/user/algorithm/index.html for algorithm details), its reference clade is assigned to the query sequence and the query sequence is phylogenetically placed relative to this node or tip. This process allows for the underlying reference phylogeny to be utilized in clade assignments, better matching the methodology used in A(H5) clade nomenclature. Nextclade also reports the query sequences’ sets of mutations, which can be useful in studying viral evolution—for instance, to identify arising variants or to understand if mammalian-adaptive substitutions are present. Further, sequence quality control scores are generated based on the aforementioned indicators—a feature useful to researchers generating and analyzing their own A(H5) HA sequence data.

## Methods

### Generation of reference phylogenies

Three reference datasets containing HPAI A(H5) HA sequences with clade annotations were obtained from the WHO/FAO/WOAH H5 Working Group: one that included sequences from each of the 55 historic HPAI A(H5) clades (n=407), one that included sequences from clade 2.3.2.1 and its descendants (2.3.2.1-like and 2.3.2.1a–g; n=2,422), and one that included sequences from clade 2.3.4.4 and its descendants (2.3.4.4-like and 2.3.4.4a–h; n=10,079). To enable assignment of LPAI lineages in addition to established HPAI A(H5) clades, we supplemented this dataset with an additional 10 Am-nonGsGd and 8 EA-nonGsGd sequences from GenBank (accessions available in **Supplementary Table S1**). We next built 3 guide trees from these reference sequences using the Nextstrain pipeline [18]. For the all-clades dataset, we used all sequences in the reference set. For the 2.3.2.1 and 2.3.4.4 datasets, we randomly sampled up to 50 sequences per clade per year and supplemented this dataset with one randomly chosen sequence from each unrepresented clade to serve as outgroup sequences. These outgrouped sequences are included to prevent erroneous Nextclade assignments to the most basal clade (i.e., 2.3.2.1 or 2.3.4.4) when query sequences from other clades are analyzed (see detailed description below). Sequences for each dataset were aligned with MAFFT [19] and divergence phylogenies were inferred using IQ-TREE 2 [20]. TreeTime [21] was used to re-root the phylogeny and to perform ancestral state reconstruction of nucleotide and amino acid sequences across every branch of the tree. The final trees contain 432 (all-clades), 1,654 (2.3.2.1), and 1,900 (2.3.4.4) sequences, including those from outgroups.

### Determination of clade-defining mutations

To assign query sequences to clades, Nextclade compares each query sequence to every tip and internal node on reference phylogeny to find the closest sequence match. To enable this for our A(H5) datasets, we next inferred clade-defining mutations (i.e., mutations that are unique to each clade) and used these sets of mutations to assign clades to both internal nodes and tips using Augur clades [22]. For each non-outgroup clade on the reference phylogeny, the last common ancestor (LCA) of all tips belonging to that clade was determined and this branch was used to define the start of the clade. Clade inheritance was established by assessing whether an LCA of one clade descended from the LCA of another clade; this allows for the clade-defining mutations of a parental clade to be inferred for its descendants. Then, sets of clade-defining mutations were determined starting at the root of the tree and working towards the tips. As an LCA was encountered, the set of mutations on that branch was assigned to its respective clade. If a mutation arose on the path to a descendant LCA and was located at the same site as a clade-defining mutation for a parental clade, it was also assigned to the descendent clade to overwrite the inferred parental mutation. For the all-clades tree, which does not include any basal, unassigned outgroup sequences, an amino acid present in the root sequence (HA 17D) was manually assigned to the basal EA-nonGsGd lineage to allow for clade assignment beginning at the root of the tree.

### Generation and benchmarking of Nextclade datasets

To build the Nextclade datasets, reference phylogenies were paired with appropriate reference HA sequences, including gene maps denoting their coding regions, to be used for alignment of query sequences and subsequent mutation calling. For the all-clades dataset, the ancestral HPAI A/Goose/Guangdong/1/1996 strain was chosen, while the candidate vaccine viruses A/duck/Vietnam/NCVD-1584/2012 and A/Astrakhan/3212/2020 were chosen for the 2.3.2.1 and 2.3.4.4 datasets, respectively [23]. In a config file for each dataset, amino acid motifs of interest were specified to allow for determination of potential N-linked glycosylation sites (PNGS) and cleavage site sequences. Cutoff values for quality control metrics were also specified here, which allow for an estimate of sequence quality based on factors such as the number of frameshifts or private mutations (mutations that separate a query sequence from its attachment site on the tree).

To benchmark our Nextclade datasets, we acquired a pre-released version (now v0.6.5) of LABEL—a widely used and well-validated command line software for A(H5) clade assignment—that includes the updated, preliminary clade splits. We then generated testing datasets using all unique A(H5) HA sequences available in GISAID and FluDB (accessed April 3, 2024; accessions available in **Supplementary Table S2**) by assigning a clade to each test sequence with both Nextclade and LABEL. For the all-clades dataset, each unique sequence that was not included in the reference dataset was included, while for the 2.3.2.1 and 2.3.4.4 datasets, each unique sequence from the appropriate clades that does not appear on the reference tree was included. These sequences were analyzed with the appropriate Nextclade dataset and assignments were compared to those from LABEL. To determine private mutation quality control values for each dataset, medians and 97.5^th^ percentiles of private mutation counts from these analyses were calculated and used as the ‘typical’ and ‘cutoff’ values, respectively. Additionally, to compare runtimes of the two tools, three randomly selected sets of 100, 1,000, and 10,000 sequences from the all-clades testing dataset were analyzed using the command-line interface for the all-clades Nextclade dataset and LABEL. For each of the three sets of sequences, runtimes were determined in duplicate and the fold-change in mean runtime (LABEL / Nextclade) was calculated. This runtime benchmarking was performed on a 2021 MacBook Pro with a 10-core M1 Pro CPU.

### Data and code availability

Accessions for the LPAI GenBank sequences used in the reference phylogenies are provided in **Supplementary Table S1** and those for the GISAID sequences used in the testing datasets are provided in **Supplementary Table S2**. Scripts and other files used in the generation of the reference phylogenies and plots are available at https://github.com/moncla-lab/h5-nextclade. Additionally, the three Nextclade datasets can be accessed at https://github.com/nextstrain/nextclade_data/tree/master/data/community/moncla-lab/iav-h5/ha.

## Results

### Generation of Nextclade datasets with A(H5)-specific features

Using WHO/FAO/WOAH-annotated reference sequence datasets—for all A(H5) clades, for clade 2.3.2.1 and its descendants, and for clade 2.3.4.4 and its descendants—we constructed three reference phylogenies. Additionally, for the 2.3.2.1 and 2.3.4.4 trees, one randomly chosen sequence from each unrepresented clade was added to serve as an outgroup sequence (to prevent erroneous downstream assignments to the most basal clade). While the all-clades dataset includes annotation of 2.3.4.4 and 2.3.2.1 subclades, using additional datasets specific to these allows for better resolution of the phylogeny for these currently circulating clades. As Nextclade compares query sequences to both tips and internal nodes of the reference phylogeny, we next determined clade-defining mutations based on the common ancestor of all tips of a given clade. These mutations were used to assign clades to internal nodes, yielding reference phylogenies with clade annotations on all tips and nodes, except for the basal outgroup sequences where applicable. Given the high circulating diversity of A(H5) viruses and the desire for these Nextclade datasets to be used on sequences from any time, quality control cutoffs for private mutations (mutations that separate a query sequence from its placement on the reference tree) had to be adjusted for these datasets. Based on analysis of unique sequences from GISAID, the median number of private mutations was used as the ‘typical’ value and the 97.5^th^ percentile as the ‘cutoff’ value (24 and 97 for all-clades; 5 and 25 for 2.3.21; 6 and 18 for 2.3.4.4) (**Supplementary Figure S1**). If private mutation counts exceed the ‘typical’ value, Nextclade will flag this QC parameter with an increasing score, with a ceiling of 100 when the value exceeds the ‘cutoff’ value.

After alignment, Nextclade will report all nucleotide mutations and amino acid substitutions relative to the dataset’s reference sequence. Using this set of mutations, the most closely related tip or node on the reference tree is determined, and the clade annotation of the tip/node is assigned to the query sequence. Further, the tool will search for N–X–S/T (where X is any amino acid except proline) amino acid motifs in the HA ectodomain of query sequences and annotate each as a PNGS. Additionally, we added in a feature to annotate amino acid sequences at sites 341–346 (all-clades) or 341–345 (2.3.2.1 and 2.3.4.4) of HA—corresponding to the polybasic cleavage site motif in each reference sequence—to produce an annotation for the cleavage site, and important determinant of pathogenicity for A(H5Nx) viruses. If four or more arginines and/or lysines are present in the motif, it is also marked as being a polybasic cleavage site.

### Dataset benchmarking and performance

To assess the accuracy of clade assignments generated by these Nextclade datasets, we utilized LABEL v0.6.5 for benchmarking. Given that this is a pre-released version and the new subclade designations are not yet finalized by the WHO/FAO/WOAH, it is possible that there are inaccuracies in its clade assignments; however, it allows us to compare assignments for any A(H5) HA sequences, including those not officially annotated, rather than a small subset of officially annotated sequences. Using all available unique HA sequences that were not found in the reference dataset, comparisons were made between Nextclade and LABEL clade assignments. The all-clades dataset performed well, with a 94.8% match rate (n=19,833) (**Figure 1A**); of note, no mismatches between HPAI A(H5) clades and LPAI A(H5) lineages were observed. Performance was best for clades with many available sequences, presumably due to better representation of within-clade diversity on the reference tree. The 2.3.2.1 and 2.3.4.4 datasets produced very well-matched assignments, with match rates of 97.8% and 99.1%, respectively (**Figure 1B, 1C**). Similarly to the all-clades dataset, these generally performed best for clades that were well represented on the reference tree. Additionally, poor resolution of the 2.3.2.1c–like and 2.3.4.4–like clades, which act as outlier groups and represent sequences that do not fall within the newly designated subclades, appears to play a large role in the mismatches seen in these datasets. Given the role of the “like” clades (i.e., to capture sequences that do not neatly cluster within a defined subclade), different approaches vary in the stringency with which sequences may be allocated into these clades, which we expect may account for many mismatches observed between LABEL and Nextclade. To compare runtimes, randomly selected sets of 100, 1,000, and 10,000 sequences were analyzed with the all-clades Nextclade dataset (via the command-line interface) and LABEL on a laptop computer, and the total execution times were determined. Nextclade very quickly analyzed large queries, performing 10,000 assignments in under 20 seconds (**Figure 2A**). With these rapid execution times, Nextclade greatly outperformed LABEL for each of the testing sets, with the runtime of LABEL being approximately 100× longer than Nextclade when 1,000 or more sequences were analyzed (**Figure 2B**).

**Figure 1.**
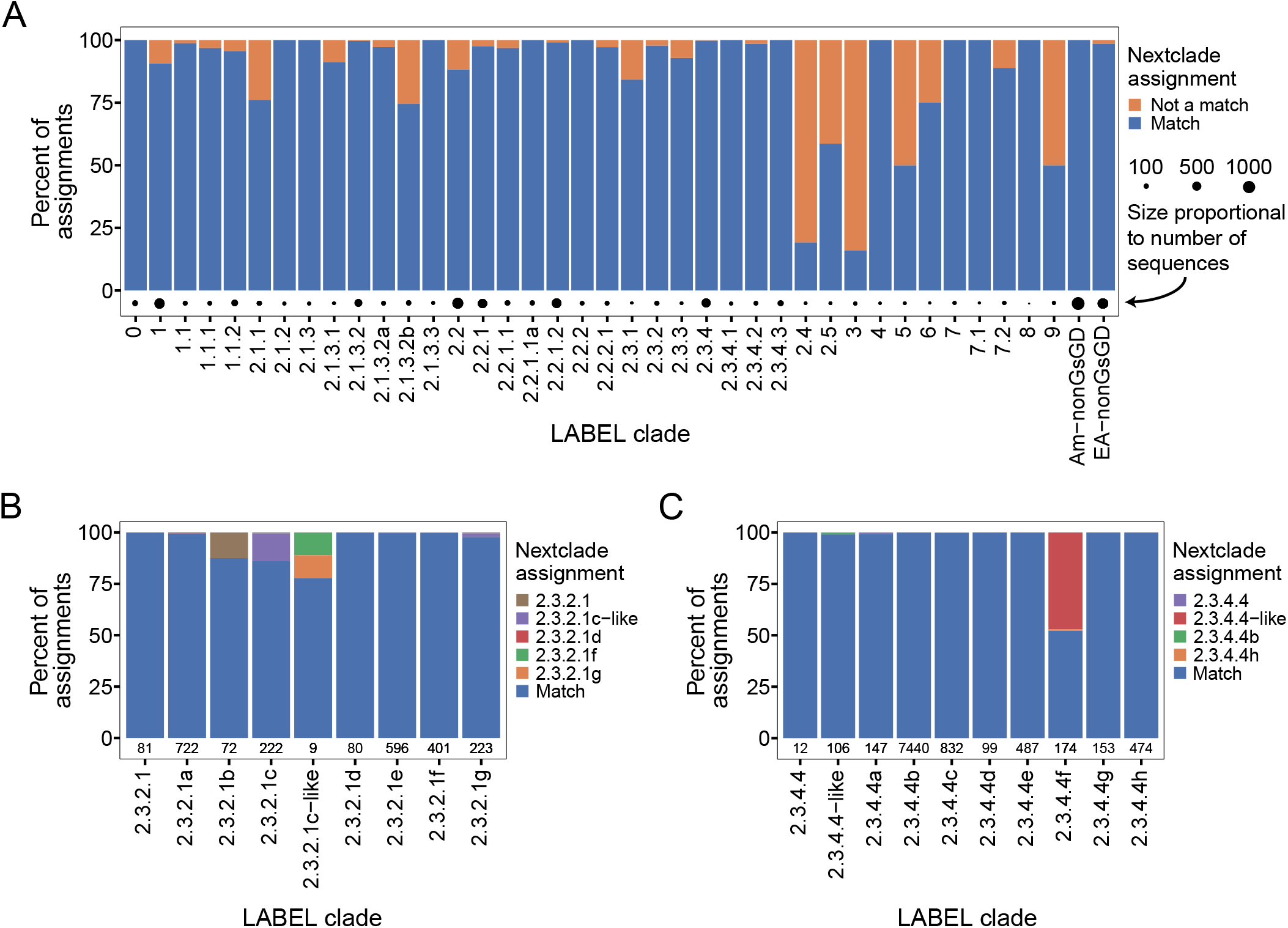
Clade assignments are well-matched between our Nextclade datasets and LABEL. Results for benchmarking against LABEL are shown for each dataset. (A) Results for the all-clades dataset (94.8% match rate, n=19,833; results for 2.3.2.1 and 2.3.4.4 descendants not shown), with blue indicating matched and orange indicating mismatched assignments. (B) Results for the clade 2.3.2.1 dataset (97.8% match rate, n=2,406), with blue indicating matched assignments and mismatched assignments shown colored by the Nextclade assignment (2.3.2.1g in orange, 2.3.2.1f in green, 2.3.2.1d in red, 2.3.2.1c–like in purple, and 2.3.2.1 in brown). (C) Results for the clade 2.3.4.4 dataset (99.1% match rate, n=9,924), with blue indicating matched assignments and mismatched assignments shown colored by the Nextclade assignment (2.3.4.4h in orange, 2.3.4.4b in green, 2.3.4.4–like in red, and 2.3.4.4 in purple).

**Figure 2.**
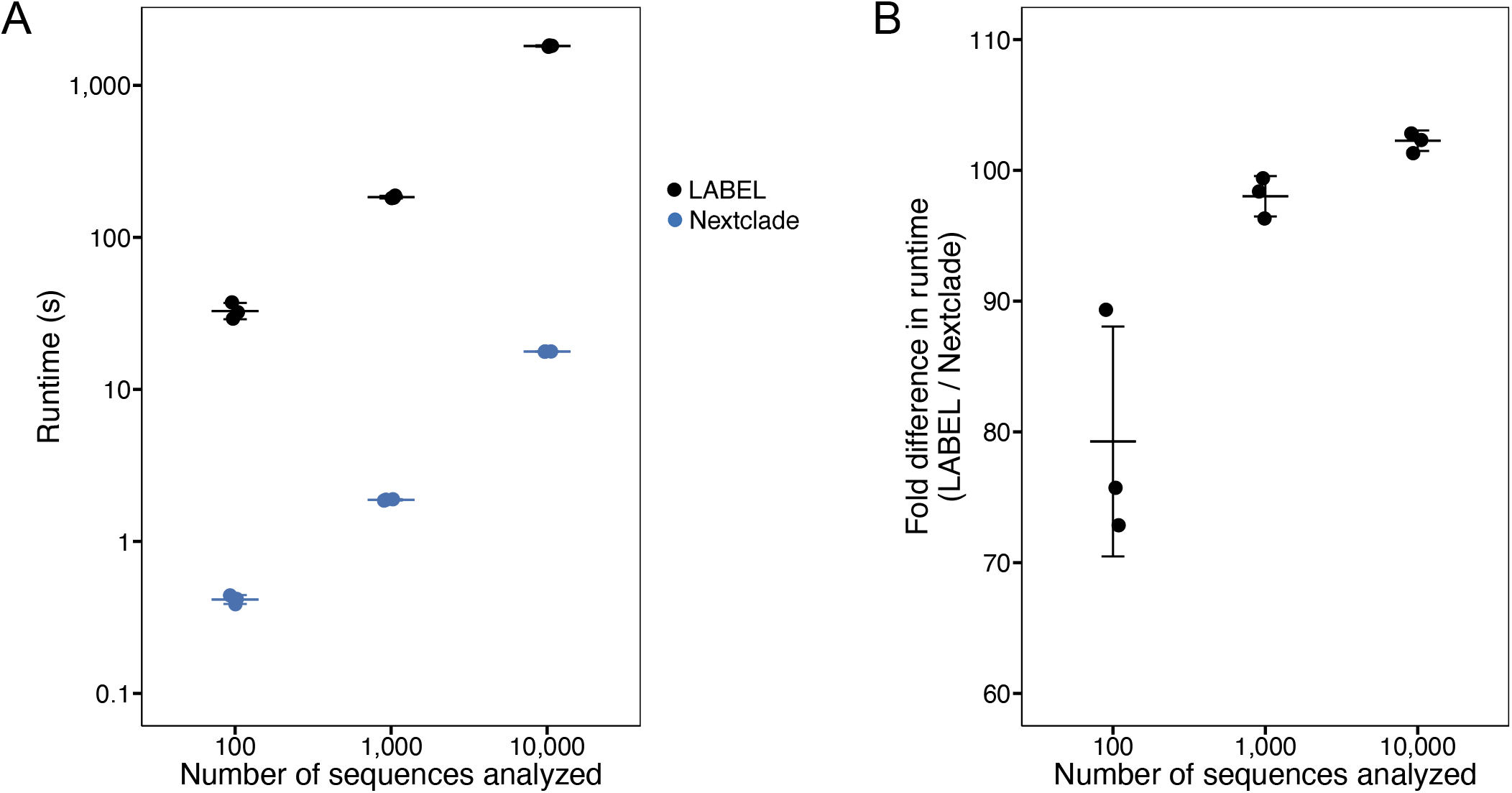
Nextclade analyzes sequences more rapidly than LABEL. Three randomly selected sets of 100, 1,000, and 10,000 sequences were annotated via the command-line interface of the all-clades Nextclade dataset and LABEL in duplicate. (A) Mean absolute runtimes were determined in seconds (s) and (B) fold differences in means were calculated, with each dot representing the value for a unique set of sequences and the mean and standard deviation shown by error bars.

## Discussion

We have generated three Nextclade datasets that can be used to assign clades to both HPAI and LPAI A(H5) sequences. Using WHO/FAO/WOAH-annotated reference sequences, we generated reference phylogenies and determined clade-defining mutations for each represented clade. The all-clades dataset can be used for all historical clades and will call mutations relative to the ancestral HPAI GsGd strain, while the 2.3.2.1 and 2.3.4.4 datasets are specific to currently circulating clade groups and will call mutations relative to candidate vaccine viruses. Nextclade will also automatically call the cleavage site sequence—and determine if a polybasic cleavage site is present—as well as locate potential N-linked glycosylation sites in the HA ectodomain. These features provide users with further details about their sequences, including those useful in identifying potential alterations to pathogenicity or antigenicity [12–14, 24, 25]. While other available clade assignment tools can provide clade assignments, we believe that Nextclade’s user-friendly interface, rapid execution, and additional features will enable a broad audience to fully leverage their A(H5) sequence data. We demonstrate that tree-based classification of avian influenza viruses is possible with Nextclade, thus future work should include the expansion of this tool to accommodate other avian influenza clade systems, such as A(H7) and A(H9). Additionally, updating these A(H5) datasets upon the designation of new clades will be critical to their continued utility. As the reference datasets can be easily amended to reflect clade updates—through the reassignment of clades to reference sequences or the addition of newly annotated sequences—Nextclade is poised to rapidly respond to nomenclature changes as they arise.

## Acknowledgements

We gratefully acknowledge labs that have generated sequence data deposited into public databases, and the WHO/FAO/WOAH H5 Working Group for continuous updates of clade designations.

