## Supplementary Figure S1 for "Development of avian influenza A(H5) virus datasets for Nextclade enables rapid and accurate clade assignment"

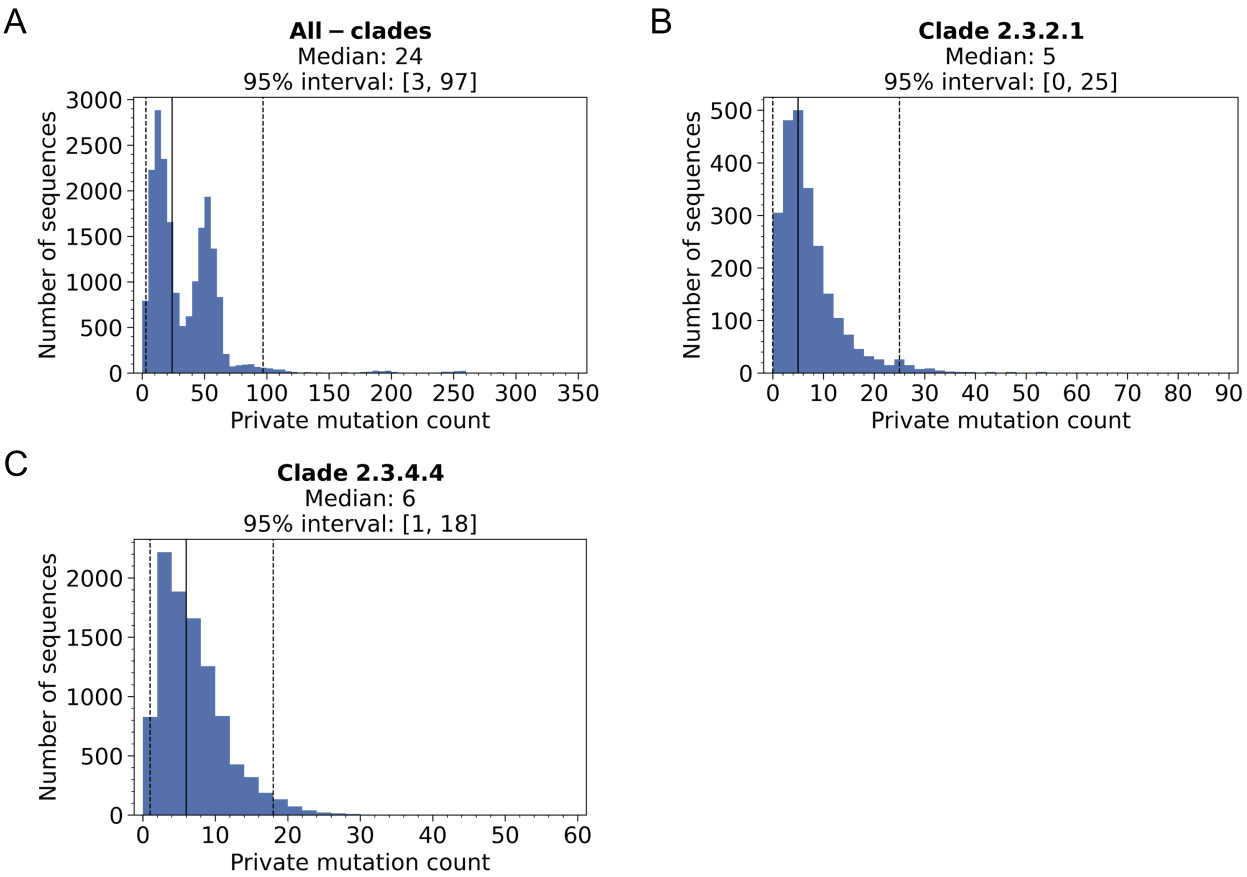


**Supplementary Figure S1. Private mutation counts for unique GISAID sequences.** Using test datasets consisting of unique sequences from GISAID that were generated for assignment benchmarking, private mutation counts of each sequence were determined and plotted as a histogram for the (A) all-clades dataset, (B) 2.3.2.1 dataset, and (C) 2.3.4.4 dataset. The median and 95% intervals are shown with a solid line and dotted lines, respectively, with both annotated above each plot.
